# Host outnumbered: microbiomes and a fungal parasite interact to shape host genotype-dependent survival

**DOI:** 10.64898/2026.01.31.703031

**Authors:** Amruta Rajarajan, Manon Coone, Justyna Wolinska, Piet Spaak, Ellen Decaestecker

## Abstract

Microbiomes are key determinants of host health, yet empirical evidence demonstrating their influence on host-parasite interactions is limited. We conducted a proof-of-principle experiment using the water flea *Daphnia magna* and its virulent yeast para-site *Australozyma monospora* (formerly *Metschnikowia bicuspidata*) to test if micro-biome composition alters infection outcomes. Axenic hosts were inoculated with a control microbiome (homogenates of laboratory-cultured *Daphnia* reared in natural freshwater) or a single bacterial strain (*Rhizobium* sp.), and exposed to the parasite. Host survival differed markedly between microbiome treatments and depended on parasite exposure. Prior to parasite exposure, hosts of one genotype exhibited high juvenile mortality when inoculated with the control microbiome (79%), compared to *Rhizobium* (19%) (the other genotype: 48% vs. 50%). Parasite exposure reduced survival, but the extent varied with microbiome composition: survival of hosts with a control microbiome (averaged across genotypes) declined from 66% to 0%; survival of those inoculated with *Rhizobium* sp. declined from 35% to 10%. In contrast, micro-biome composition did not influence parasite infectivity or net reproduction. Our results indicate that microbiome effects on host survival are genotype- and life-stage-dependent, differ between parasite-free and parasite-exposed conditions and may impact host-parasite dynamics primarily by influencing host viability rather than para-site performance.

## Introduction

Symbioses between hosts and microbes are ubiquitous in nature, and span a continuum from mutualistic to parasitic interactions. Host-associated microbial communities (microbiomes) are integral to host health and are linked to disease (Hou et al., 2022; Izvekova, 2022), and are increasingly consideredin wildlife conservation and disease ecology (Trevelline et al., 2019). Microbiomes can influence hosts’ immune-related gene expression across diverse taxa; including fish (Koch et al., 2018), plants (reviewed in Hacquard et al., 2025) and insects (reviewed in Schmidt & Engel, 2021), and can affect pathogen colonization resistance through resource competition (Spragge et al., 2023). Observational studies in natural systems, such as amphibian chytridiomycosis (Jani et al., 2017) or epidemics of ichthyosporean gut parasite in water fleas belonging to the *Daphnia longispina* complex (Rajarajan et al., 2025) reveal correlations between microbiome profiles and disease epidemics *in situ*. Moreover, the presence of mutualistic symbionts can alter host-parasite co-evolutionary trajectories. (Rafaluk-Mohr et al., 2022).These findings underscore the need for experimental approaches to establish ecological pathways linking microbiome compositions and disease outcomes.

Host genetic background and microbiomes are both linked to host survival and may contribute to the heterogeneity in susceptible host populations shaping disease dynamics (Rose et al., 2021). Early-life microbiomes in human beings are hypothesized to interact with genetic background to influence longevity and disease (Kadyan et al., 2025). Microbiome-driven effects on host survival are especially challenging to quantify because of the inherent survivor bias in microbiome surveys (Lampeter et al., 2023). Nevertheless, one rigorous investigation revealed that gut microbiome composition correlate with future survival of bird hosts across breeding seasons *in situ*, although the specific effect of host genetic background is unknown (Worsley et al., 2021). In laboratory-based studies, cross-species microbiome transplants reduced survival in the snail hosts *Biomphalaria* (Schols et al., 2023), dung beetle *Onthophagus* (Parker et al., 2021), as well as broadly in sister-species pairs across vertebrate and invertebrate hosts (Brooks et al., 2016) suggesting that survival is species-specific and mediated by microbiome compositions.

Microbiome-transplant approaches are vital to establish causal connections between hosts and their health and disease outcomes, yet highlight a fundamental challenge: complex microbiome transplants do not constitute identical treatments across genotypes. Host genotypes widely associate with distinct microbiomes (Ahern et al., 2021; Callens et al., 2018; Stagaman et al., 2017). In germ-free hosts, genetic background can influence priority effects in host-colonization dynamics of transplanted microbiota (Gurung et al., 2024; Leopold & Busby, 2020) or differentially filter transplanted microbiomes through immune regulation and further influence the function of microbiota (reviewed in Wilde et al., 2024). As a result, differences in host survival following microbiome transplantation may reflect not only the inoculated microbiota themselves, but also their genotype-specific assembly by the host. Thus, to understand if host genotypes differ in their microbiome-driven responses, it is essential to transplant individual strains across host genotypes, alongside complex microbial communities reflecting ecological reality (Greyson-Gaito et al., 2020).

To fully understand ecological pathways shaping disease dynamics, it is necessary to examine causal relationships between host genetic background and microbiome compositions on one hand, with host survival and parasite fitness on the other. Here we used a microbiome-transplant approach with two genotypes of the freshwater zooplankter *Daphnia magna* susceptible to the yeast parasite *Australozyma monospora* to test the effect of microbiome composition on the outcome of the host-parasite interactions. Germ-free hosts were inoculated either with a single bacterial strain (*Rhizobium* sp.) or with microbiomes derived from donor individuals reared in natural lake water, and subsequently exposed to the parasite. We asked whether microbiome composition: (1) alters host survival in the presence and absence of parasite exposure, (2) affects parasite infectivity and net reproduction, and (3) does so consistently across host genotypes.

## Material & Methods

### Host-parasite system (Daphnia magna and Australozyma monospora)

Complex microbiome- and genotype-mediated effects on host survival may be especially prevalent in aquatic environments where microbiomes are transient and strongly influenced by environmental conditions (Eckert et al., 2021; Russell, 2019; Stock et al., 2021), resulting in dynamic symbiont associations and functions (Gillingham et al., 2025).The freshwater zooplankter *Daphnia magna* is a key model organism in host-parasite-microbiome investigations due to its clonal reproduction and widespread ecological occurrence (Ebert, 2022, Macke et al. 2017). Microbial community composition within individual *Daphnia* hosts varies widely *in situ*, ranging from a few strains to complex and rich communities (Houwenhuyse et al., 2025; Rajarajan et al., 2025).

The yeast parasite *Australozyma monospora* (formerly *Metschnikowia bicuspidata*, taxonomy revised by (Lachance et al., 2025) is a commonly occurring generalist parasite termed an ‘obligate killer’, i.e. infectious particles can only be transmitted after host death (Stewart Merrill & Cáceres, 2018). Infection occurs through ingestion of waterborne spores during filter feeding. After penetrating the gut epithelium, the parasite proliferates in the hemolymph, ultimately causing high host mortality and releasing infectious spores into the environment (Stewart Merrill & Cáceres, 2018). *Australozyma* exposure reduces microbial alpha diversity and alters bacterial community composition in a genotype-specific manner, regardless of the establishment of an infection, suggesting that parasite-driven microbiome effects may influence host tolerance (Rajarajan et al., 2024).

#### Microbiome transplant procedure

The experimental procedure involved transplanting either control microbiome inocula consisting of pooled-individual *Daphnia magna* homogenates (one per genotype E-17-07 and NO-V-7), or a single strain of *Rhizobium*, to germ-free recipients. Bacterial community compositions of control inocula were characterized with 16S rRNA metabarcoding. Due to the previously reported microbiome differences between the genotypes E-17-07 and NO-V-7 (Rajarajan et al., 2024), control inocula were only transplanted to recipients of the same genotype, resulting in four groups: two host genotypes × two microbiome treatments. Microbiome-inoculated recipients were then experimentally exposed to the parasite *Australozyma monospora* to compare host (juvenile mortality, adult survival) and parasite traits (infectivity and net reproduction) between genotypes and microbiome treatments.

To generate control inocula, juveniles born within 24 h were reared individually in 20 mL medium consisting of supplemented natural microbiota (see supplementary material) and fed *C. vulgaris* (5 ×10^5^ frozen cells/ mL) three times per week. Fresh medium was added as needed to compensate for evaporation. Offspring produced during this period were discarded within 24 hours. At 15-16 days of age, *Daphnia* were transferred individually with a pair of forceps to 1.5 mL microcentrifuge tubes containing 300 µL 0.22 µm-filtered, autoclaved tap water and homogenised. Homogenates were pooled by host genotype to create control inocula. These inocula were filtered (0.7 µm) to remove debris, stored at 4 °C, and used within 8 h. An aliquot of each control inoculum (one per genotype) was metabarcoded following Rajarajan et al., (2024).

Axenic microbiome recipients were created following Callens et al., (2018). To generate individuals with eggs at the optimal developmental stage (12-24 hours after deposition in the brood pouch), recipient cultures were maintained as described for stock cultures (see supplementary material). Briefly, eggs were removed from brood pouches using sterilised forceps, washed with 0.1% glutaraldehyde (v/v) for 10 s, rinsed twice and transferred to a parafilm-sealed six well plate containing 0.22 µm-filtered, autoclaved tap water and incubated at 20°C, 16:8 light:dark cycle for 72 hours until hatching.

Free-swimming juveniles were pooled by genotype and randomly assigned to microbiome treatments. Recipients were placed individually in 25 mL sterile-filtered, autoclaved tap water and inoculated with either: (i) their genotype-specific “control” inoculum at a donor:recipient ratio of 1:2, or (ii) a 48-h old *Rhizobium sp*. culture (“monoculture”). The monoculture inoculum dose was standardized to control inocula based on OD_600_ measurements (Mushegian et al., 2016). All individuals remained in their microbiome treatments (n = 42 per genotype × microbiome group) for 72 h prior to parasite or placebo exposure.

#### Parasite/placebo exposure

24 hours before parasite exposure, all individuals received 2 × 10^6^ cells/mL autoclaved *C. vulgaris*. Within each microbiome treatment, half of the surviving individuals were assigned to the parasite exposure treatment. Microbiome-inoculated hosts, together with 5 mL of their medium, were transferred to 15 mL falcon tubes and exposed to either 3,500 *Australozyma* spores/mL or an equivalent volume of placebo solution. After 48 h, individuals were transferred back to their original tubes in which they received control inocula, while minimizing transfer of spore-containing medium, and topped up to 40 mL sterile-filtered autoclaved tap water. Recipients were subsequently maintained at 20 °C (16:8h light:dark), and fed autoclaved *C. vulgaris* (6 × 10^6^ cells/mL) three times per week.

#### Recipient host and parasite life-history data collection

Juvenile survival was recorded after the 72-h microbiome inoculation period. Thereafter, host survival was monitored daily. Dead individuals were fixed in 3% formaldehyde within 24 h of death. Beginning 10 days post-exposure, live individuals were screened for *Australozyma* infection under a stereomicroscope (20×), without being removed from the tube. The experiment was terminated when the last infected individual died (day 14 post-exposure). Fixed specimens were homogenized with a pestle, and parasite spores were quantified using a Neubauer chamber (mean of three technical replicates), regardless of treatment.

#### Statistical analyses

Statistical analyses were conducted in R (R Core Team, 2023) for host traits (juvenile survival, survival post parasite- or placebo exposure) and parasite traits (infectivity, parasite transmission). Data was filtered with dplyr (Wickham H et al., 2023), data.table (Barrett T et al., 2025), vegan (Dixon, 2003), ape (Paradis & Schliep, 2019) and microbiome (Lahti & Shetty, 2017) R packages and visualised using ggplot2 (Wickham, 2016) and survminer R (Kassambara A et al., 2025) packages. Due to low reproduction in germ-free animals re-inoculated with microbiomes, host fecundity was not analysed.

*Juvenile survival* (binary survival-outcome following the 72-h microbiome inoculation) was analysed using χ2-tests. Because control inocula differed between genotypes (see Results), a full-factorial model would not disentangle genotype from microbiome effects. We therefore performed three χ^2^-tests on contingency tables: (i) comparing juvenile mortality between genotypes within the *Rhizobium-*monoculture treatment, and (ii) comparing microbiome treatments separately for each genotype. Tests were performed with *chi*.*sq* function in base R.

*Host survival* after parasite or placebo exposure was analysed with Cox proportional hazards models. Individuals that died prior to parasite exposure were excluded. First, we fitted a mixed-effects model: survival probability ∼ microbiome * parasite exposure + (1 | genotype) using survival (Therneau, 2023) and coxme (Therneau, 2025). Host genotype was modelled as a random effect because of unbalanced sample sizes between genotypes due to juvenile mortality in the E-17-07 control microbiome group (see Results). As genotype explained negligible variance (0.008%), we adopted a simpler model structure: survival probability ∼ microbiome * parasite exposure, for which sample sizes were: control-placebo n=14, monoculture-placebo n=26, control-parasite n=16, monoculture-parasite n=30. Cox models were fitted using the *coxph* function from the survival R package, with pooled data across genotypes.

*Parasite infectivity* was calculated as the proportion of successfully infected individuals (based on presence/absence of mature *Australozyma* asci upon death) among those exposed to the parasite and alive at least 10 days post-exposure, as infection status can only be assessed after this timepoint (Stewart Merrill & Cáceres, 2018). Infectivity was analysed using a General Linear Model (GLM) with the structure: infection outcome ∼ microbiome * genotype with the binomial family distribution using the lme4 R (Bates et al., 2024) package.

*Net reproduction* of the parasite was defined as the number of spores produced per exposed host, with uninfected or early-dead individuals assigned a value of zero to capture total contribution of spores to the next generation. Net reproduction was modelled as a continuous variable using a General Additive Model (GAM) and the mgcv (Wood, 2025) R package. GAM structure was parasite transmission ∼ micro-biome * genotype with a Tweedie distribution.

## Results

### Control inocula of the two genotypes differed in dominant bacterial genera

Control-microbiome inocula consisted of 26 dominant bacterial genera (Fig. 1). These 26 most abundant genera accounted for 84% of the bacterial community in genotype E-17-07 and 61% in genotype NO-V-7 (Fig. 1). Most dominant genera were geno-type-specific, except *Limnohabitans* (4% in E-17-07 and 20% in NO-V-7) and *Flavobacterium* (4% and 3%, respectively) which were abundant in control inocula of both genotypes. *Rhizobium* was not detected in any control-microbiome inoculum. The most abundant genera were *Lactococcus* in E-17-07 (45%) and *Limnohabitans* in NO-V-7 (20%). Overall, control inocula contained 114 genera in E-17-07 and 112 in NO-V-7, of which 100 genera were shared between genotypes.

**Fig. 1.**
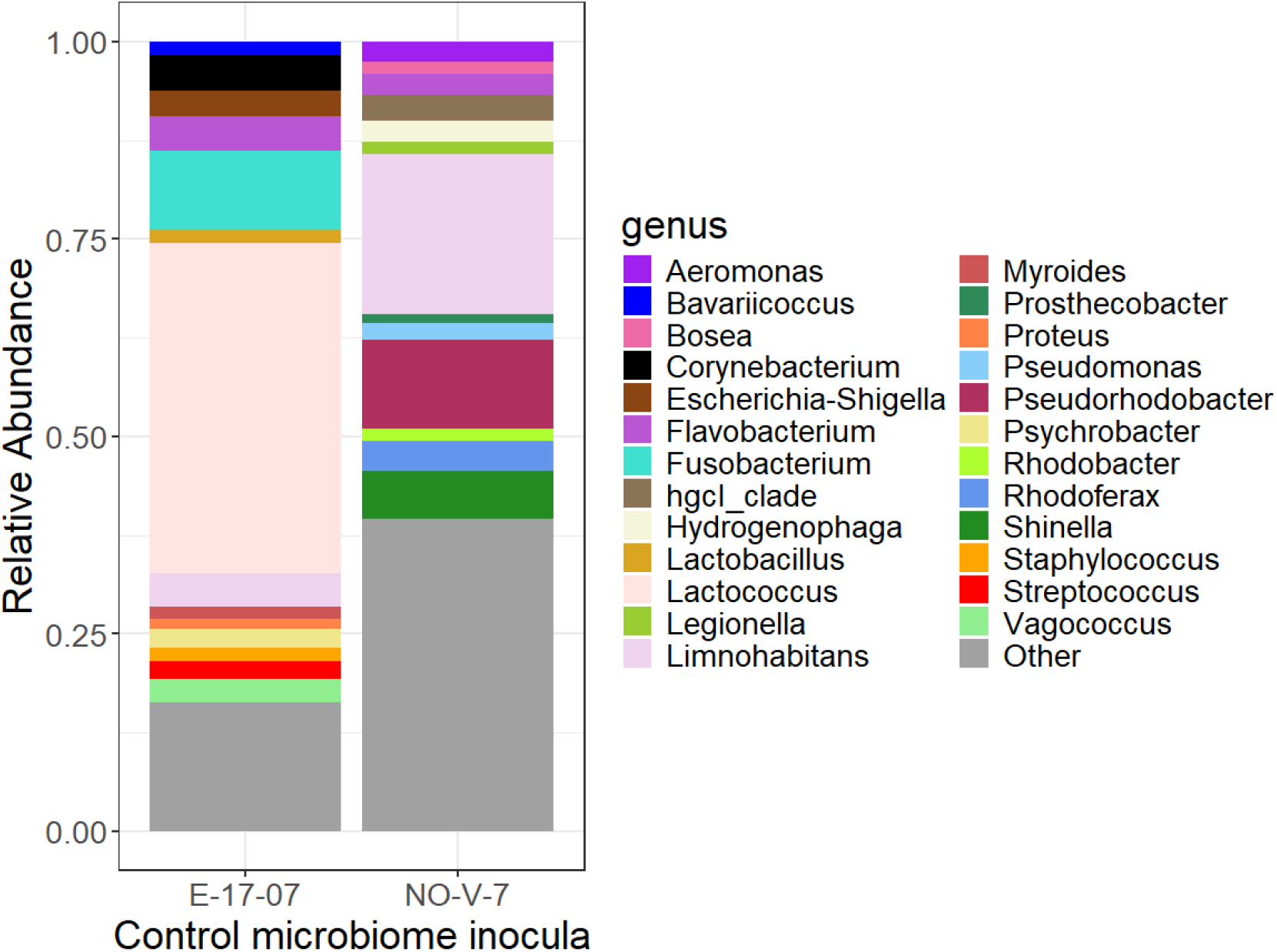
Relative abundances of dominant bacterial genera in control microbiome inocula for each host genotype (genotypes indicated on the x-axis). The 26 most abundant bacterial genera in control inocula are indicated with colours and all other genera are grouped as „Other” (grey). Each inoculum represents pooled whole-individual homogenates of 15 adult *D. magna*.

### Juvenile survival was highest in E-17-07 hosts inoculated with *Rhizobium*

Juvenile survival after the 72-h inoculation period and prior to parasite exposure differed between microbiome treatments in genotype E-17-07 but not NO-V-7 (Fig. 2). In E-17-07, survival was significantly lower in individuals receiving the control microbiome (19%) than in those inoculated with *Rhizobium* sp. (79%, χ^2^ = 27.5, *p* < 0.001). In contrast, juvenile survival in NO-V-7 did not differ between microbiome treatments (48% vs. 50%; χ^2^ = 0, *p* = 1). Comparing genotypes within the *Rhizobium* treatment, E-17-07 juveniles exhibited lower survival than NO-V-7 (χ^2^ = 6.27, *p* = 0.01). These results show that both microbiome composition and host genotype shape juvenile survival.

**Fig. 2.**
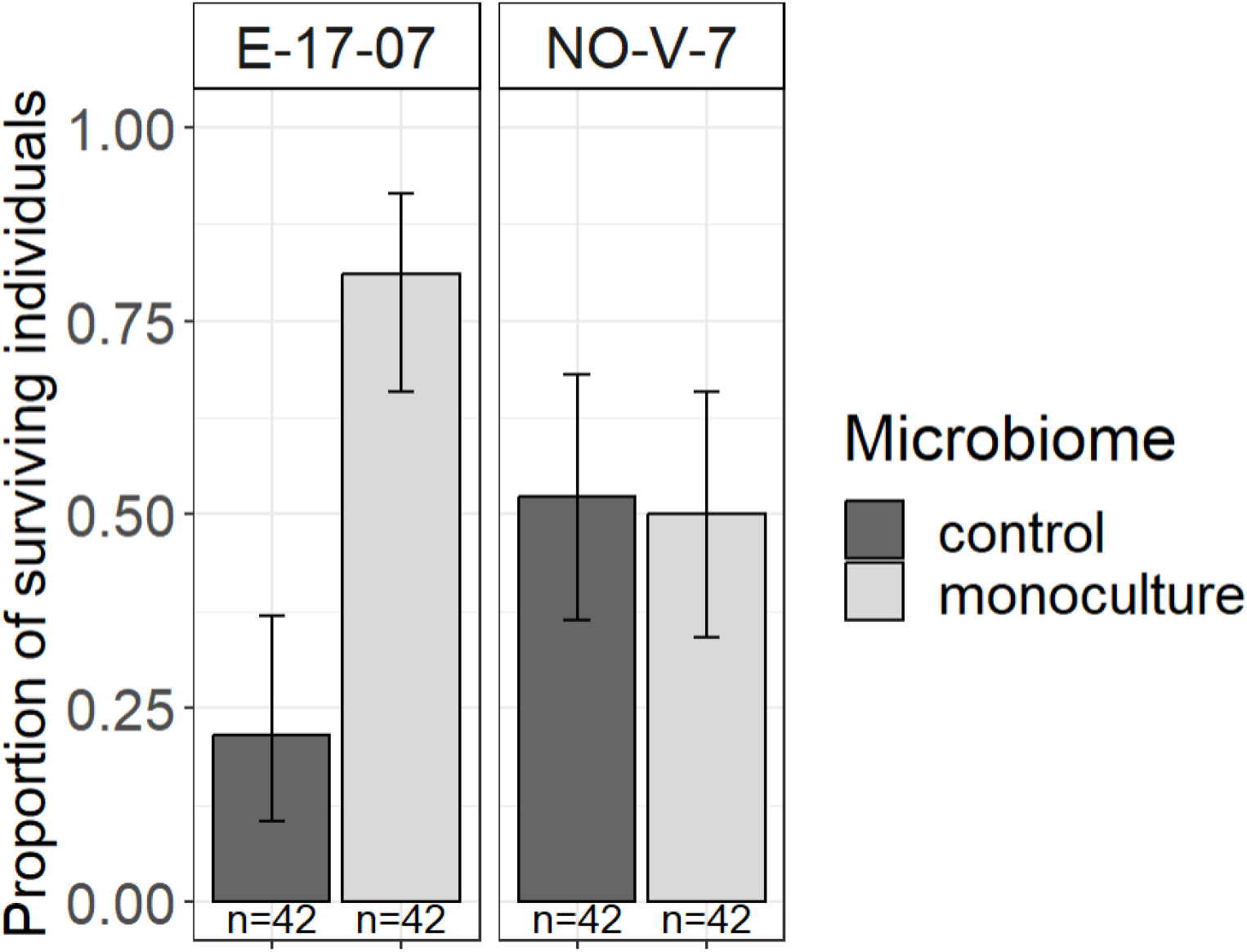
Proportion of juveniles surviving the 72-hour microbiome inoculation period prior to parasite exposure (juvenile survival). Host genotypes (E-17-07 and NO-V-7) are separated into panels, and colours indicate microbiome treatment. Whiskers denote confidence intervals estimated using a binomial distribution. Microbiome composition did not affect NO-V-7 juvenile survival. E-17-07 juvenile survival was significantly higher in the monoculture treatment compared to control. The monoculture significantly increased survival in E-17-07 compared to NO-V-7.

### Host survival varies marginally with the interaction between parasite exposure and microbiome composition

Parasite exposure significantly reduced host survival in both microbiome treatments (Fig. 3, z = -4.1, *p* < 0.001, Cox proportional hazards). Microbiome treatment alone had no effect (z = -1.03, *p* = 0.3), but the microbiome × parasite interaction was marginally significant (z = 1.88, *p* = 0.06). Specifically, hosts with a control microbiome experienced a greater survival cost of parasite exposure, with survival declining from 66% in the placebo treatment to 0% following parasite exposure. In contrast, *Rhizobium*-inoculated hosts exhibited a smaller reduction in survival, declining from 35% to 10%.

**Fig. 3.**
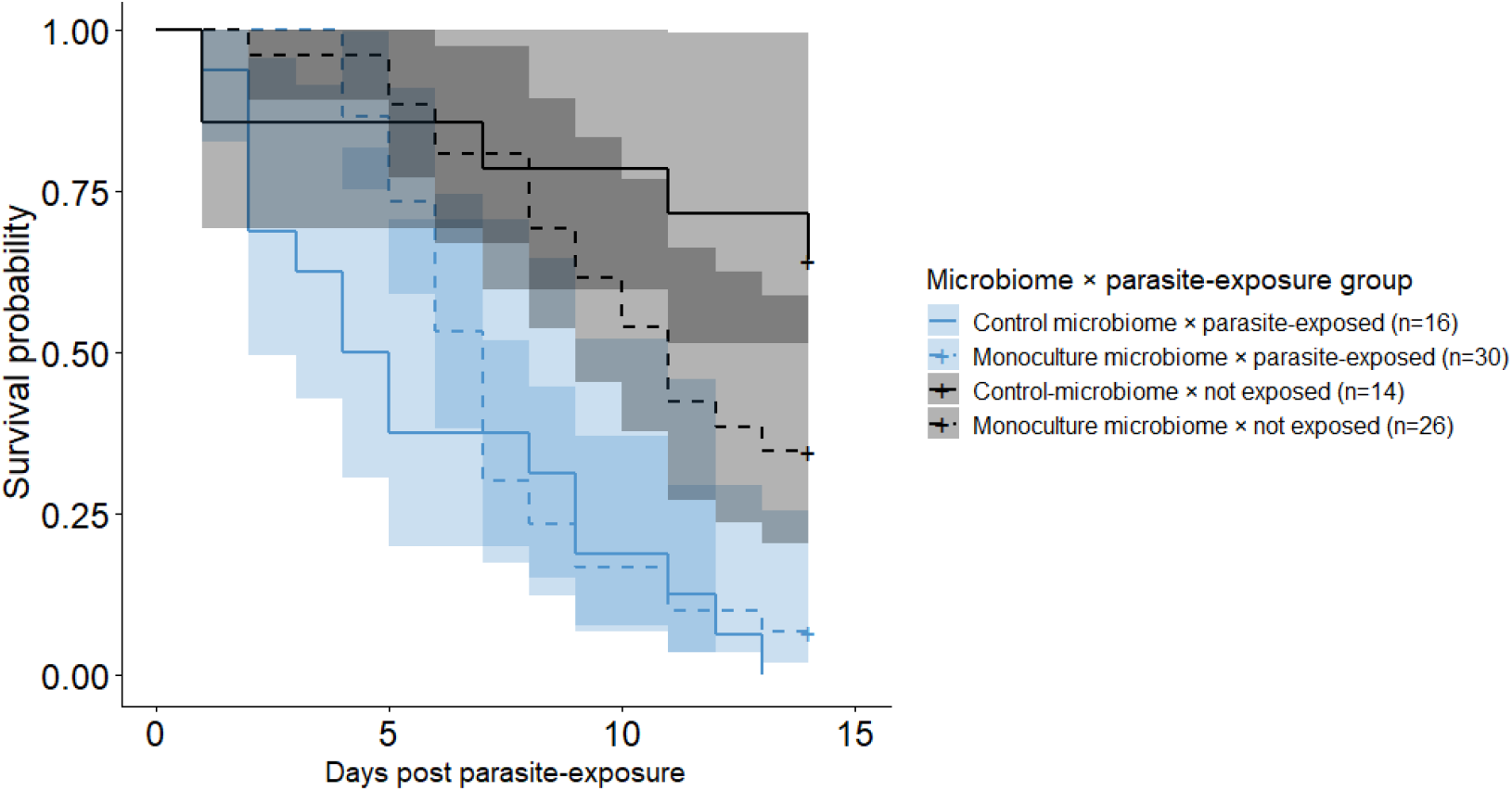
Survival probabilities of hosts over time for *D. magna* individuals inoculated with control microbiomes (solid lines) or *Rhizobium* monocultures (dashed lines) in the placebo (black) and parasite-exposed treatments (blue). Host genotypes are pooled within each treatment. Shaded areas denote 95% confidence intervals. *Autralozyma* exposure significantly reduced host survival in both microbiome treatments, while microbiome treatment had no effect. Only individuals that survived the 72 h microbiome inoculation are included; juvenile survival was disproportionately low in genotype E-17-07 inoculated with the control microbiome prior to parasite exposure (see Fig. 2).

### Parasite infectivity and net reproduction did not differ by host genotype or microbiome treatment

Parasite infectivity did not differ between microbiome treatments, genotypes, or their interaction (Fig. 4A). In the control microbiome group, infectivity was 40% (infection probability; 95% CI: 10-80%) in E-17-07 and 37 (95% CI: 14-66%)in NO-V-7 (z = - 1.4, *p* = 0.89, GLM). In the *Rhizobium* group, infectivity was slightly higher (48% [95% CI: 27-69%] and 47% [95% CI: 20-73%], respectively), but this difference was not statistically significant (z = 0.29, *p* = 0.77, GLM).

**Fig. 4.**
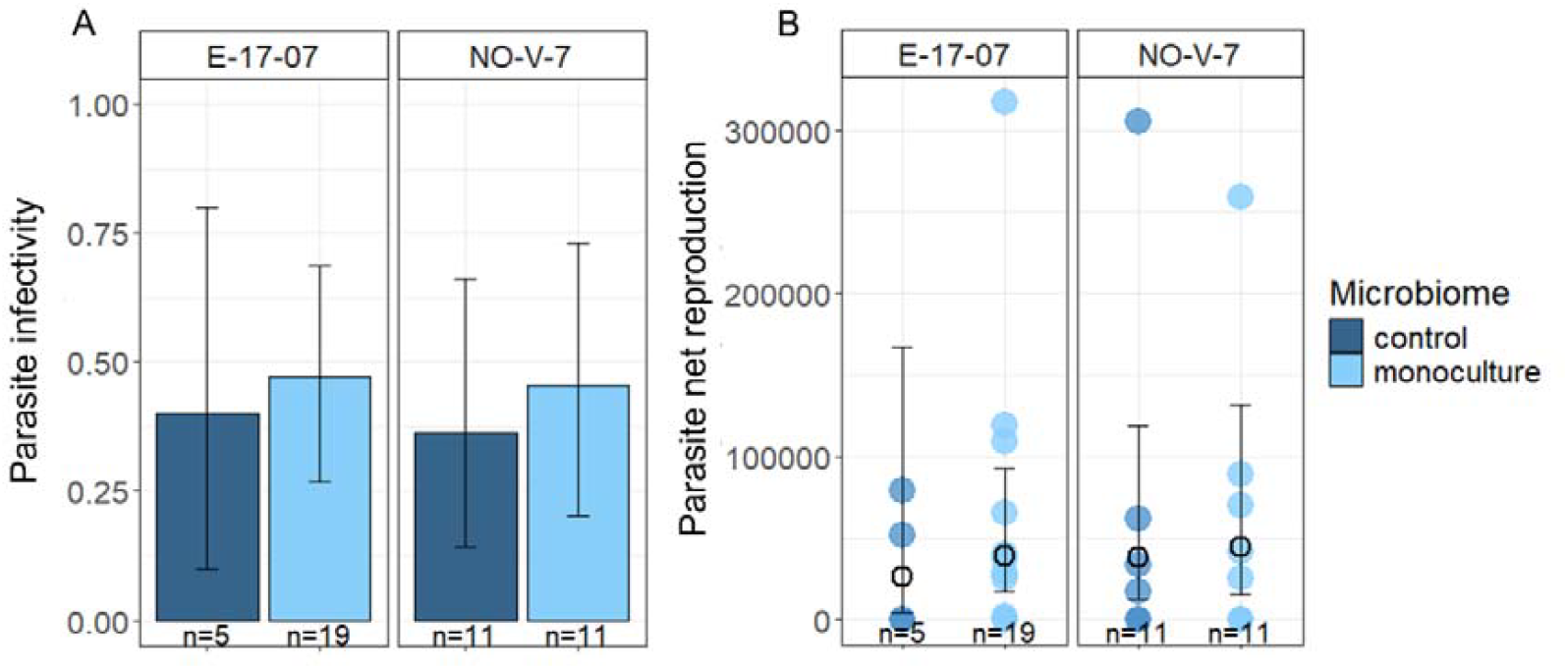
Parasite infectivity (A) and net reproduction (B) in host genotypes E-17-07 and NO-V-7, with microbiome treatment indicated by colour. (A) Infectivity was calculated as the proportion of successfully infected individuals among those exposed to the parasite. Whiskers indicate confidence intervals estimated with a binomial distribution. (B) Parasite net reproduction, quantified as the number of spores produced per exposed host. Points represent individual hosts, with colours denoting microbiome treatment. Black symbols indicate mean values ± confidence intervals estimated with a Tweedie distribution. Individuals exposed to the parasite that did not get infected (i.e. produced 0 spores), either due to failure of infection or host inviability, are included.

Parasite net reproduction (spores produced per exposed host) likewise did not differ between microbiome treatments or host genotypes (Fig. 4B). Genotype E-17-07 produced on average 26,000 spores (95% CI: 0-165,000) in the control microbiome group and 38,947 (95% CI: 0-90,000) spores in the *Rhizobium* group (t = 0.38, *p* = 0.7, GAM). Genotype NO-V-7 showed similar patterns (37,878 [95% CI: 0-120,000] vs. 43,939 [95% CI: 0-130,000]spores; t = 0.34, *p* = 0.74).

## Discussion

Host microbiome compositions are frequently correlated with disease, yet empirical tests of their causal effects on disease outcomes remain scarce. Here we used a microbiome-transplant approach to test whether microbiome composition influences the outcome of a host-parasite interaction by affecting host (*Daphnia magna*) and parasite (*Australozyma monospora*) fitness traits. Overall, we found that the impact of microbiome composition on juvenile survival was host genotype-specific, but that parasite traits were not directly affected by microbiome composition. This suggests that microbiome effects on host-parasite interactions may arise primarily through indirect effects on host viability rather than through direct modulation of parasite performance.

### Host genotype and microbiome compositions interact to influence host survival

In our study, microbiome composition did not affect juvenile survival in genotype NO- V-7, whereas survival was strongly reduced in genotype E-17-07 – unexpectedly when hosts were inoculated with microbiomes originating from donor individuals of the same genotype reared in control conditions. Methodological artefacts arising from the germ-free procedure are unlikely to explain this result, since eggs of both genotypes were made germ-free simultaneously using a standardised procedure (see Methods) whereas only one genotype showed strong microbiome-dependent mortality; and bacterial (see Table S1) or eukaryotic pathogen-driven mortality was not observed in laboratory-reared animals. Moreover, the same *Rhizobium* strain had a significantly positive effect on juvenile survival of one genotype (E-17-07) compared to the other (NO-V-7), suggesting that overall result is consistent with a general geno-type-microbiome interaction effect on host survival (Coone et al., 2023; Houwenhuyse et al., 2021).

We speculate two mechanisms by which microbiomes derived from healthy individuals may cause mortality when transplanted to germ-free individuals of the same genotype. First, the relatively low abundance of *Limnohabitans*, a keystone microbe known to enhance *Daphnia* reproduction and survival (Bisschop et al., 2025; Gurung et al., 2026; Peerakietkhajorn et al., 2015) in E-17-07 hosts (see Fig. 1A) may explain why mortality is higher in this genotype compared to NO-V-7. Second, the juvenile mortality may have resulted from a potential mismatch of life stages across which the microbiome transplant was conducted – i.e. from adult donors to juvenile recipients.

Similar age-specific microbiome effects were reported in *Drosophila* hosts: *Acetobacter*, a symbiont commonly found in association with adults (Kim et al., 2016) shortened lifespan when inoculated into juvenile *Drosophila* (Obata et al., 2018). Our results would further suggest that a hypothesized critical developmental window for microbiome composition (Fontaine & Trevelline, 2025) potentially differs between genotypes. Future studies could incorporate wider ranges of genotypes and microbiome compositions to quantify genotype-specific, microbiome-induced juvenile mortality is *in situ*. Overall, our findings suggest that specific genotype- and life-stage-specificity must be considered when implementing microbiome-based interventions for wildlife population health.

Regardless of the specific mechanism, demographic models predict that relatively small shifts in survival may have consequences at the population level (Barrus et al., 2025). For example, copper-induced mortality – particularly in juveniles – played a key role in reducing population growth of the freshwater rotifer *Brachionus calyciflorus* (Schanz et al., 2021). Our results therefore suggest that natural variation in microbiome compositions has the potential to shape host population dynamics by disproportionally affecting the survival of particular genotypes or life stages. Evolutionary theory further predicts that such microbiome-induced changes in host fitness may alter host evolutionary trajectories (Henry et al., 2021). Future studies could explicitly link microbiome-induced survival differences to population dynamics in host-parasite systems.

### Microbiome composition does not impact parasite traits but may influence host-parasite dynamics

Although microbiomes can influence *D. magna* nutrition (Callens et al., 2016) and thus potentially the within-host environment experienced by the parasite, we found no evidence that host microbiome compositions affected *Australozyma*’s infectivity or net reproduction. Despite the absence of direct effects on parasite traits, our results suggest that microbiome compositions may impact disease dynamics by influencing genotype-specific early life mortality. *Daphnia* microbiome beta diversity co-varies with the population density of hosts, as well as with the size of natural disease epidemics (Rajarajan et al., 2025). Moreover, the outcome of *Australozyma* infection is strongly host genotype-dependent (Duffy & Sivars-Becker, 2007). Here we show that microbiome composition induced significant juvenile mortality in one of the two genotypes susceptible to the parasite. This raises two ecological pathways by which natural variation in microbiome compositions could influence parasite epidemics: (i) by reducing the number of competent hosts available for infection, and (ii) by altering the genetic composition of the host population available to the parasite.

### Microbiome-driven host survival is context-dependent and may not depend on complexity

The relationship between microbiome richness and host health is debated;symbioses between hosts and their microbes may have complex and context-dependent effects on disease ecology due to the ability of symbionts to shift along a mutualism-parasitism continuum (Hopkins et al., 2017). A more diverse microbiome is often proposed to offer colonization resistance against parasites, for example due to more effective nutrient blocking (Spragge et al., 2023). Aligning with this hypothesis, a richer microbiome was associated with increased tolerance to *Ranavirus* in amphibian hosts (Harrison et al., 2019) and increased resistance to *Flavobacterium columnare* in the zebrafish host (Stressmann et al., 2021). Alternatively, a richer microbiome may not confer defence against a parasite if the increased richness is primarily of “non-core” microbial taxa, as shown in bumble bee hosts (Cariveau & Winfree, 2015).

Consistent with complexity-specific microbiome function, co-infection by parasites can trigger symbiont shifts along the mutualism-parasitism continuum. In separate experiments, co-infection with the mildly virulent microsporidian parasites created a net benefit to *Daphnia* hosts in the presence of more virulent parasites, such as the microsporidium *Hamiltosporidium tvaerminnensis* (O’Keeffe et al., 2024), White Fat Cell Disease-causing virus (Lange et al., 2014) and *Australozyma* (Rogalski et al., 2021). Our study revealed a trend suggesting the same microbial strain (*Rhizobium*) may be mildly deleterious in parasite-free conditions while being mildly beneficial in the presence of a parasite, whereas complex microbiomes had a significantly negative genotype- and age-specific impact on host survival. Therefore, the identity, or interactions of key symbionts - including parasites - with specific host life stages, rather than microbiome richness or complexity *per se* may determine survival.

## Conclusion

Our results demonstrate that host microbiome composition may impact disease dynamics by shaping genotype-specific juvenile mortality and thus altering the demography of host populations exposed to the parasite. Importantly, microbiome compositions did not influence parasite infectivity or transmission, indicating that microbiomes may not directly mediate resistance. We also found a trend suggesting that the same microbial strain (*Rhizobium*) may provide mild benefits to parasite-exposed hosts while being mildly deleterious in parasite-free conditions, implying potential context-dependent symbiont functions. Collectively, these findings suggest that host micro-biomes may influence disease epidemics primarily by altering host viability.

## Acknowledgement

We thank the Disease Evolutionary Ecology group (IGB Berlin) for providing *Daphnia magna* genotypes and the *in vivo* parasite strain as well as the Aquatic Biology group (KU Leuven KULAK) and Spaak groups (Eawag) for discussions on experiment design and encouraging feedback. We thank the Genetic Diversity Center (GDC) at ETH Zürich for supporting metabarcoding library preparation and bioinformatic processing for metabarcoding data. We also thank Aditi Gurung and Shinjini Mukherjee (KU Leuven) for isolating the *Daphnia*-associated *Rhizobium* strain and Isabel Vanoverberghe for extensive experimental support. We are grateful to two anonymous referees for their constructive criticism on the manuscript. This work was funded by KU Leuven research proposal C16/17/002, Eawag (Swiss Federal Institute of Aquatic Science and Technology) and a joint “lead agency” grant from the German Science Foundation (WO 1587/6-1 to JW) and Swiss National Science Foundation (310030 L 166628 to PS).

## Author contribution statement

**Amruta Rajarajan:** Conceptualisation (equal); Methodology (equal); Formal investigation (lead); Writing - original draft (equal); Writing - review & editing (lead). **Manon Coone:** Methodology (equal); Formal investigation (supporting); Data curation (lead); Writing - original draft (equal); Writing - review & editing (supporting). **Justyna Wolinska:** Conceptualisation (equal); Formal investigation (supporting); Writing - review & editing (supporting); Funding acquisition (equal); Supervision (equal). **Piet Spaak** Conceptualisation (equal); Funding acquisition (equal); Supervision (equal). **Ellen Decaestecker**: Conceptualisation (equal); Methodology (supporting); Formal investigation (supporting); Writing - review & editing (equal); Funding acquisition (equal); Supervision (equal).

