## Supplementary material for "Host outnumbered: microbiomes and a fungal parasite interact to shape host genotype-dependent survival"

**Table S1**. Results of Web of Science literature searches conducted on 5^th^ May 2026 to discover evidence of pathogenicity among genera detected in *Daphnia* microbiomes. Each search was conducted with the terms *[genus name] AND Daphnia AND (pathogen* OR parasit*)* where [genus name] corresponds to the genera reported as dominant in the E-17-07 control inoculum, and absent in NO-V-7 control inoculum. Abstracts of hits were screened to find evidence of pathogenicity towards *Daphnia*. While the bacterial genera have generally been reported to occur in association with *Daphnia*, no evidence for pathogenicity was found.

| **Genus name** | **Number of hits** |
| --- | --- |
| *Bavariicoccus* | 0 |
| *Corynebacterium* | 2 |
| *Escherichia-Shigella* | 4 |
| *Fusobacterium* | 14 |
| *Lactococcus* | 11 |
| *Myroides* | 2 |
| *Proteus* | 11 |
| *Psychrobacter* | 3 |
| *Staphylococcus* | 16 |
| *Streptococcus* | 13 |
| *Vagococcus* | 14 |

General culturing conditions

**Host *Daphnia* *magna*:** Two *Daphnia magna* genotypes were used: E-17-07 (England) and NO-V-7 (Norway). Stock cultures were maintained at 20 °C with a 16:8h light:dark regime, with 10-12 females per genotype kept in 2 L medium composed of 50% 0.45 μm-filtered Greifensee lake water (Switzerland; N 47°20′41″, E 8°40′21″) and 50% 0.22 μm-filtered tap water. *Daphnia* were fed frozen *Chlorella vulgaris* (5 × 10^6^ cells/mL), three times per week, and medium was refreshed every three weeks.

**Parasite *Australozyma monospora*:** The parasite strain “METS_AMME_2008”, isolated in 2008 from lake Ammersee (Germany), was maintained *in vivo* on the *D. magna* genotype E-17-07 following procedures described in Manzi et al., (2022). To prepare the parasite inoculum, *Australozyma*-infected hosts were homogenised with a pestle in sterile-filtered tap water; to prepare the placebo inoculum, uninfected E17:07 host*s* were similarly homogenized. Homogenates were filtered through autoclaved 0.7 µm GF/F filters (Whatman Cat #28497-914), allowing bacteria to pass while retaining spores, which were rinsed and resuspended in autoclaved and 0.22 µm-filtered tap water to avoid carryover of microbiota (Rajarajan et al., 2024). The same procedure was followed for the placebo solution. *Australozyma* spore concentration in the parasite inoculum was then quantified from two subsamples using a Neubauer chamber (Merck #MDH-4N1).

***Rhizobium* symbiont**: Members of Rhizobiales, including *Rhizobium*, have been detected in aquatic microbiomes and reported in association with *Daphnia* (Houwenhuyse et al., 2025; Rajarajan et al., 2025). *Rhizobium* species can colonize *Daphnia* guts and influence host life history traits (Gurung et al., 2024). Beyond their association with aquatic hosts, *Rhizobium* spp. are well-established mutualists in terrestrial ecosystems, contributing to host nutrition, immune modulation and pathogen resistance (Poole et al., 2018). *Rhizobium* sp. strain *R2_7*, originally isolated from *D. magna* gut tissue (Gurung et al., 2024), was revived from -80 °C glycerol stock and grown in Reasoner’s 2A medium (Reasoner & Geldreich, 1985). Ten microliters of stock were inoculated into 5 mL of Reasoner’s 2A medium under sterile conditions. A negative-control flask received an empty sterile pipette tip. Cultures were incubated at 25 ˚C, with shaking at 125 rpm until turbid and were transferred every three days (10 µL into 10 mL fresh Reasoner’s 2A medium). Negative controls were maintained in parallel throughout to validate the absence of contamination.
